# Selective Glucocorticoid Receptor Modulators of Immune Checkpoint Function

**DOI:** 10.64898/2026.02.02.703046

**Authors:** Robin R. Kobylski, Charles K. Min, Jerome C. Nwachukwu, Marcel Koenig, Haotian Zhang, Lizbeth Moncada, Thomas T. Venables, Bin-Bin Schell, William J. Stein, Claudia McCown, Justin Schumate, Emelie Bonilla, Amin Sobh, Arantxa Romero-Toledo, Matthew E. Pipkin, Michalina Janiszewska, Louis Scampavia, Timothy P. Spicer, Jonathan D. Licht, Inez Rogatsky, Theodore M. Kamenecka, Laura A. Solt, Kendall W. Nettles

## Abstract

Glucocorticoids (GCs) coordinate immunity, inflammation, and metabolism through allosteric regulation of the glucocorticoid receptor (GR) transcription factor. GCs are indispensable anti-inflammatory drugs yet linking specific ligand–receptor structural states to specific biological outcomes has remained a major barrier to designing safer, more selective therapies. Using structure-based design, we developed selective glucocorticoid receptor modulators (SGRMs) of immune function by extending a steroidal scaffold from the ligand-binding pocket into an adjacent solvent channel. These SGRMs suppressed T cell pro-inflammatory cytokines and promoted differentiation of memory precursor T cells while showing minimal induction of M2 macrophage polarization or T cell checkpoint proteins PD-1 and CTLA-4, all key targets of immunotherapy. Molecular dynamics simulations revealed that solvent-channel substituents function as a lever arm to drive dynamic oscillations in the steroid core, thereby allosterically tuning GR activity states. Systematic perturbation of immune cells with a graded series of ligands enabled a *l*igand *<u>p</u>*erturbation with *<u>m</u>*achine *l*earning (LPML) framework to map coregulated responses across cell types and identified effector T cell gene networks tightly coupled with immune checkpoint induction. This approach outlines a general strategy for decoding the logic of allosteric drug action, enabling the rational design of SGRMs with tailored immunomodulatory profiles.

## Introduction

Ligand-regulated allosteric receptors such as glucocorticoid receptor (GR) can adopt distinct structural states that drive different biological outcomes, termed selective modulation (*1*). GR serves as a primary organismal stress response system, coordinating transcriptional programs that respond to infection, injury, inflammation, and nutrient deprivation (*2-4*). Endogenous cortisol and synthetic glucocorticoid (GCs) such as dexamethasone (Dex) are widely used anti-inflammatory agents, but their broad actions can impair host defense and antitumor immunity and promote metabolic liabilities including hyperglycemia, obesity, osteoporosis, and muscle wasting (*5*). Despite extensive molecular and clinical characterization, the organizing principles that couple GR structural states to tissue-, cell type-, and pathway-selective outcomes remain poorly defined, constraining rational development of selective GR modulators (SGRMs).

This dilemma stems from the very nature of GR’s mechanism as a ligand-regulated receptor as it can adopt a large ensemble of structural states, each capable of driving distinct biological outcomes. Perturbing cell states with a single ligand yields large molecular datasets—including GR interactions with transcription coregulator complexes, genomic DNA binding profiles, and regulated genes—but does not readily reveal the dynamic causal connections that define primary mediators of specific biological outcomes (*6, 7*) (**Fig. 1A**). To address this, we generated closely related ligands that target the same structural feature of GR, enabling *repeated ligand perturbation* of a specific allosteric pathway with systematically varied signals. This structured variance focuses analytical attention on features such as cellular responses that rise and fall in concert with defined phenotypes and allows quantification of coordinately regulated activities across pathways, cell types, and tissues using machine learning, revealing the underlying circuitry. Within this repeated *l*igand *p*erturbation with *m*achine *l*earning (LPML) framework, recurrent patterns of coregulation identify networks that couple ligand structure and receptor conformation to cellular and organismal responses.

**Figure 1.**
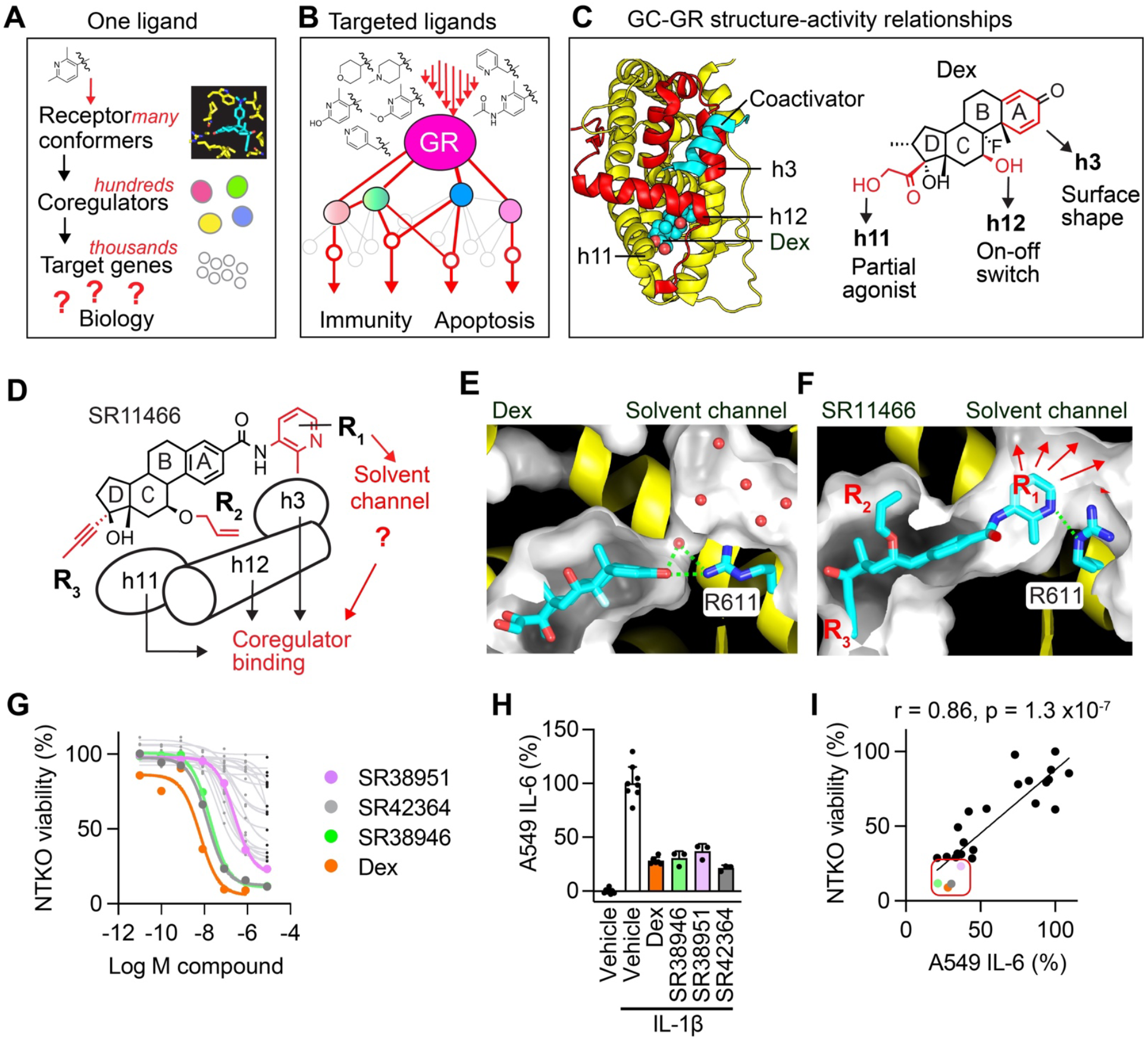
A structure-based design strategy yields potent modulators that target the GR solvent channel. **(A)** Challenges in identifying causal links between GCs and their effects. See also **Fig. S1A. (B)** Designing a series of ligands to target a specific structural feature identifies specific causal pathways through patterns established by repeated ligand perturbation. **(C)** Crystal structure of GR ligand binding domain with a peptide from the SRC-2/Grip-1 coactivator peptide (cyan) with Dex shown as spheres. Right, chemical structure of Dex with steroid rings lettered A-D. Different allosteric signaling paths are indicated. **(D)** Chemically addressable structure-based design based on SR11466, a muscle-sparing SGRM. **(E)** Cartoon of Dex-bound GR crystal structure showing the ligand binding cavity and solvent channel, connected by a small pore with ordered water (red spheres) linking the two sites. **(F)** Molecular dynamics simulation of SR11466-bound GR with R_1_ methylpyridine extended into the solvent channel. **(G)** Effects of R_1_ targeted GC ligands on viability of NTKO multiple myeloma cells. N = 8. **(H)** IL-6 secretion by A549 lung carcinoma cells treated with IL-1β and GCs as indicated. N = 4. **(I)** LPML correlation plot showing the relationship between the effects of the R_1_ modified GCs on NTKO cell viability and IL-6 secretion by A549 cells. Each dot represents the maximum effects of a distinct compound. Compounds are colored as in panels **G-H**. Pearson correlation, r, calculated from SGRM activity excluding Dex and vehicle.

Previous applications of LPML identified estrogen receptor modulators with efficacy in models of endometriosis and endocrine-resistant breast cancer as well as GCs with muscle-sparing properties (*8-10*). When assessing GCs, ligand-based computational analysis revealed transcriptional coregulator–gene networks in skeletal muscle that control glucose disposal and glucocorticoid-mediated protein balance, providing a path to rationally designed SGRMs (*9*). Here, we have extended this strategy to immune regulation by GR and discovered SGRMs with selective immune modulatory properties, linking a steroid scaffold to chemical moieties that bound in the solvent channel adjacent to the conserved ligand binding site. Our computational approach identified a network of GC regulated genes that highly correlated with induction of the immune checkpoint receptors PD-1 and CTLA-4. Further analysis revealed coordinate regulation between core GR signaling networks and genes implicated in T-cell function, memory, metabolism, and exhaustion. Allosteric regulation from the solvent channel was chemically addressable for specificity to other steroid receptors, suggesting a conserved and modifiable allosteric mechanism that can be more generally used to engineer pathway-selective modulators. Collectively, our approach offers an opportunity to tune anti-inflammatory and checkpoint-related activities while preserving essential host-protective functions.

## Results

### Targeting the GR solvent channel for directed ligand perturbation yielded potent modulators of receptor signaling

By developing a set of ligands with a range of activities that target a specific structural feature, we generated structured variance in their biological effects as tools to reveal patterns of coregulated signaling across GR target cell-types and pathways (**Fig. 1B)**. We developed a strategy to target different allosteric signaling mechanisms with chemically addressable modifications to the base steroid agonist scaffold at specific locations (**Fig. 1C–D**). These mechanisms included targeting the known agonist/antagonist switch comprised of the c-terminal helix 12 in the ligand binding domain, which directly regulates the shape of the binding site for transcription coregulators (**fig. S1A–C**). The binding events for coregulator complexes to specific genomic loci are the primary mechanisms for transcriptional regulation by nuclear receptors, including an ensemble of histone modifying enzymes that regulate chromatin structure and polymerase activity (*4, 11*). We also identified indirect mechanisms of allosteric regulation (*8, 12-14*) that produced selective biological outcomes for ER and GR ligands (*9, 10, 15*). Guided by these design principles, we synthesized a series of SR11466-derived ligands with modifications at each of the indicated targeting sites on the steroid scaffold with R_1_ modifications extending into the solvent channel, a conserved region in the steroid receptors that lies underneath the coactivator binding surface and opens from the surface to a small pore in the ligand binding pocket of steroid receptors (**Fig. 1D**). While GR ligands such as Dex bind via hydrogen bonds to R611, Q570, and an ordered water at the interface between the ligand binding cavity and solvent channel, we found that SR11466 opened the pore to enable potent binding, previously seen with a full agonist, deacylcortivazol (*16*) (**Fig. 1E-F, fig. S1B**). SR11466 is a muscle sparing anti-inflammatory SGRM (*9*), prompting us to explore solvent channel targeting as a general mechanism to modulate receptor activity.

The solvent channel-targeted SGRMs displayed picomolar (pM) to nanomolar (nM) potency for GR in gene reporter assays (**fig. S1C, Table S1)** and spanned a full range of activities suppressing growth of non-translocated knockout cells (NTKO), a genetically modified human multiple myeloma cell line containing the *t*(4;14) translocated allele, a cancer for which GCs are a primary treatment (*17*). NTKO cells were steroid sensitive, while triple knock out (TKO) cells, lacking the translocation, were resistant to Dex and other GCs including SGRMs, demonstrating GR signaling specificity (**Fig. 1G, fig. S2A–B**). In a benchmark assay for GR anti-inflammatory activity, when compared to Dex, the best compounds, SR38946, SR38951, and SR42364, robustly inhibited secretion of the pro-inflammatory cytokine IL-6 from A549 lung cells **(Fig. 1H)**, an NF-κB dependent cytokine driver of asthma, which also promotes metastatic growth of multiple myeloma in bone (*18, 19*). Using Pearson correlation, we found that the R_1_-focused SGRMs inhibited IL-6 and NTKO viability similarly across compounds (**Fig 1I**). We also observed a similar pattern with the compound effects on inhibiting viability of *in vitro* derived CD8^+^ memory-like precursor T cells (Tmp) (*20, 21*) (discussed further below), and on C2C12 myoblasts, a mouse skeletal muscle cell line that can differentiate into myotubes under specific cell culture conditions (**fig. S2C**). In this context, growth inhibition may relate to the promotion of myotube fusion under appropriate conditions (*9*). In contrast, the correlation with effects on IL-6 and Tmp viability were weak with the steroid resistant TKO cells (**Fig. S2D**). The tight coregulation of anti-cancer growth inhibition and anti-inflammatory cytokine suppression across the gradient of ligand activity suggests these effects are driven by common GR-mediated signaling mechanisms. To determine if this pathway was conserved in primary immune cells and to probe for selective effects, we profiled the SGRMs in a panel of T cell and macrophage subsets.

### SGRMs selectively modulate immune cell activity

GCs are well-known immunosuppressants but play complex roles in immune cell differentiation that have not been fully elucidated (*22*). GCs govern thymic T cell development and modulate immune responses in the tumor microenvironment; GC biosynthesis and GR levels are coordinated across tissue cell types to control differentiation and immunity (*2, 23*), including promotion of T cell memory and recall (*20, 24*). Chronic GC exposure can lead to persistent yet dysfunctional T cells, a state known as exhaustion (*25*), and while this phenotype is associated with chronic infection and tumor progression, the gene networks underlying these diverse phenomena are not well known. Macrophage polarization to an M2 phenotype is differentially regulated by GCs or other cytokines like IL-4 (*3*), but the degree to which these effects can be modulated by SGRMs is also unclear.

We established an *in vitro* compound profiling platform to study the functional effects of targeting the GR solvent channel on immune cells. We used primary mouse immune cells to probe distinct facets of the immune response in CD4^+^ and CD8^+^ T cells and macrophages (**Fig. S3A–C**) (*3, 26, 27*). We adapted established protocols for generating CD8^+^ T cell subsets, specifically using defined concentrations of the cytokine IL-2 to determine CD8+ T cell fates *in vitro*. High IL-2 concentrations (100U/mL) were used to drive cytotoxic effector-like T cells (Teff) whereas low IL-2 concentrations (10U/mL) promoted development of Tmp (*27*). For CD4^+^ T cells, we implemented differentiation protocols previously described for generating T helper type 1 (Th1) and Th17 subsets (*26, 28, 29*) to capture the plasticity and context-dependent roles of T helper cells. For macrophages, we assessed M2-like macrophages, or alternatively activated macrophages, those that developed in the presence of the cytokine IL-4, promote anti-inflammatory responses, tissue repair, wound healing, and pro-tumoral activities (*3, 30, 31*).

When profiling CD8^+^ T cells, we found that the most efficacious SGRMs, SR38946, SR38951, and SR42364, acted similarly to Dex as full agonists in certain contexts, fully inhibiting production of tumoricidal TNFα from CD8^+^ Teff cells. The ligands also affected CD8^+^ Tmp differentiation, evidenced by increased IL-7 receptor (IL-7R) and decreased CD62L expression (**Fig. 2A**). IL-7R and CD62L are critical cell surface markers defining CD8^+^ T cell memory states and are important for regulation of survival, persistence, and lymphoid organ/tissue homing (*24, 32*). These data suggest the SGRMs can maintain host-preserving aspects of GC function through maintenance of immune memory, supporting GC regulation of a memory phenotype.

**Figure 2.**
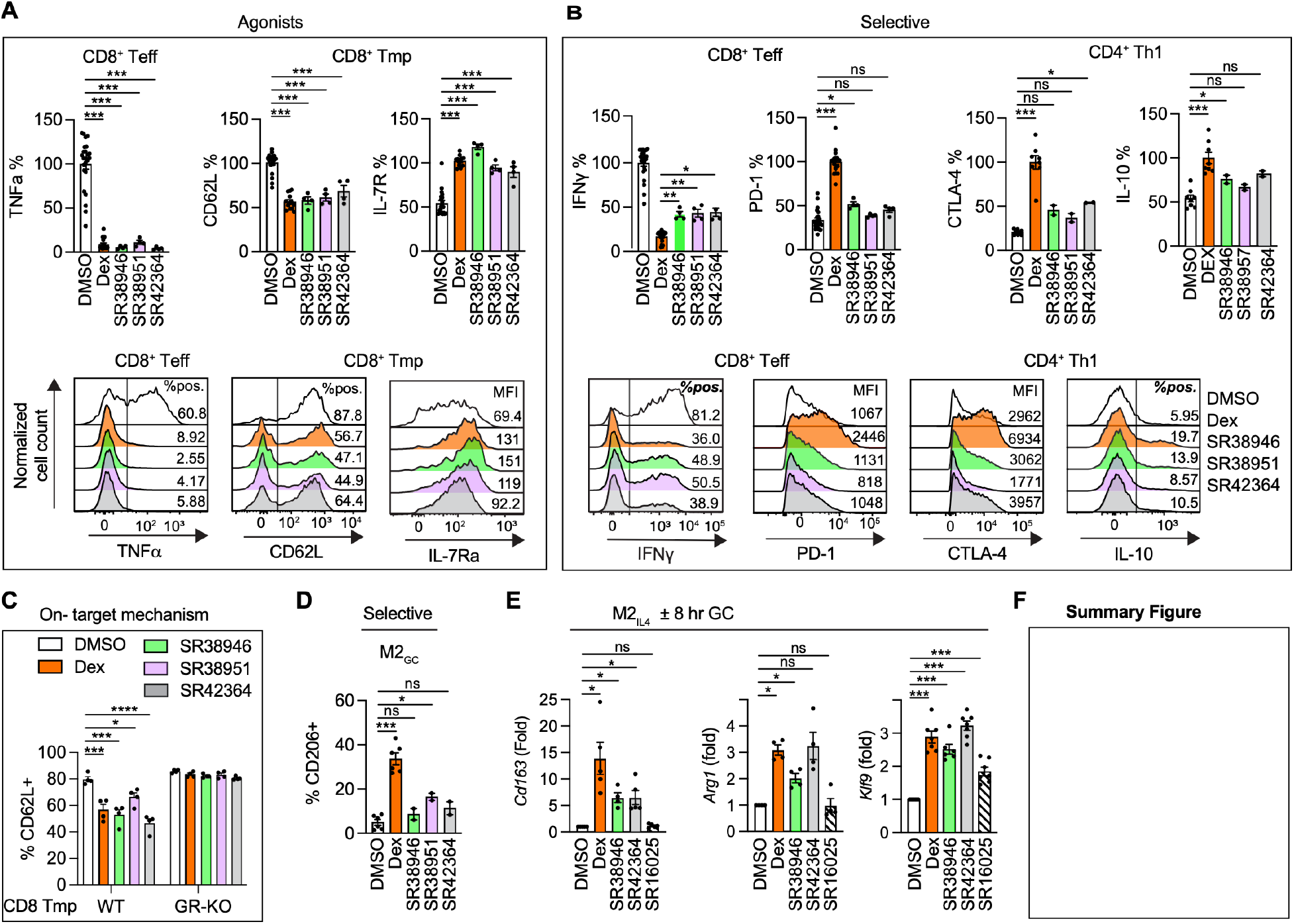
SGRMs display immune function selectivity. **(A)** TNFα, CD62L, and IL-7R expression in CD8^+^ T effector-like (Teff) and CD8+ T memory-like precursor (Tmp) cells, respectively, treated with the indicated GCs. (**B)** IFNγ, PD-1, CTLA-4, and IL-10 expression in CD8^+^ Tmp and CD4+ Th1 cells, respectively, treated with the indicated GCs. **(A–B)** Graphs (*top*) represent data normalized to vehicle or Dex treatments. n = 4-16, Mean + SEM. (*Bottom*) Representative histograms of flow cytometry data collected from each respective cell type. See also **fig. S3–4. (C)** CD62L expression in CD8^+^ Tmp cells from WT (*Nr3c1*^*+/+*^ *x dLCKCre+*) or GR knock-out (KO) cells (*Nr3c1*^*fl/fl*^ *x dLCKCre+*) treated with the indicated GCs. **(D)** Percentage of anti-inflammatory (CD11c^-^/CD206^+^) M2 macrophages differentiated in the presence of the indicated GCs. **(E)** qRT-PCR analysis of IL-4 polarized M2 cells treated with the indicated GCs. Mean + SEM of N = 4–8. Significance determined by one-way ANOVA with Dunnett’s multiple comparisons test (* p < 0.033, ** p < 0.022, *** p < 0.001).

In contrast to the Dex-like effects observed in Figure 2A, the SGRMs only partially inhibited Teff production of IFNγ, retaining significantly more IFNγ expression than the near-complete inhibition by Dex (**Fig. 2B**). This effect was also observed with IFNγ expression in Th1 cells, retaining approximately 40-50% of the stimulation induced levels relative to Dex (**Fig. 2B, fig. S4A**). The SGRMs also induced limited expression of PD-1 and CTLA-4 in CD4^+^ and CD8^+^ T cells (**Fig. 2B**), primary targets of immunotherapy that naturally limit autoimmunity and are well known aspects of GC immune suppression (*25, 33*). Production of the anti-inflammatory cytokine IL-10 by Th1 cells was also attenuated with the SGRMs (**Fig. 2B**), a counterbalancing regulator of the pro-inflammatory effects of IFNγ on the inflammatory state of the Th1 cells and microenvironment (*34*). In this context, the strong anti-inflammatory effects on Teff cell TNFα but retention of significant Teff and Th1 IFNγ expression is important, as IFNγ is required for tumoricidal T cell effects, including responses to immunotherapy (*35*). In contrast, tumors can evade TNFα induced cell death through various mechanisms, including loss of TNF/TNF receptor signaling components (*36*). As GCs are the primary therapy for adverse events downstream of PD-1 and CTLA-4 blockade, the development of SGRMs that maintain the anti-tumor efficacy of immunotherapy has important therapeutic implications. To confirm specificity of the SGRMs, GR expression was selectively ablated in post-thymic mature T cells by crossing GR floxed (Nr3c1^fl/fl^) mice (*37*) with dLCK-Cre transgenic mice (*38*). SGRM suppression of CD62L was abrogated in GR-deficient T cells, as were the known effects of GCs on Tmp cell viability (**Fig. 2C, fig. S4B**)(*20*), confirming a GR-dependent mechanism of action.

While profiling macrophages stimulated with Dex (M2_GC_), we found that, consistent with previously published results, Dex, promoted development of this subset toward an M2-like phenotype as expected, evidenced by increased expression of the M2 marker CD206 (*3*)(**Fig. 2D**). In contrast, the SGRMs did not have this effect, failing to induce this pro-tumoral state. Further, adding Dex and SGRMs to macrophages polarized with IL-4 (M2_IL-4_) demonstrated different outcomes on *Cd163, Arg1 and Klf9* gene expression, demonstrating the plasticity of macrophage differentiation to different stimuli (*3*) (**Fig. 2E, fig. S4C**). *Cd163* and *Arg1* showed variable induction by the SGRMs compared to Dex (**Fig. 2E**), genes associated with poor outcomes in tumor infiltrating macrophages (*3, 39*). SR16025, an R_2_ directed antagonist (*9*), had reduced gene activation in this context, suggesting that the different R groups can be used to develop new combinations of effects profiles (*data not shown*). This provides additional support that the solvent channel-targeting mechanism elicits broad differences in immunomodulation compared to standard GCs. These effects include maintenance of beneficial effects on T cell differentiation and cytokine inhibition while demonstrating dissociative effects on immune suppressive markers and tumor-promoting markers in T cells and anti-inflammatory macrophages (**Fig. 2F**). These selective activities define a novel and advantageous immunomodulatory profile, one that retains potent anti-inflammatory and memory-shaping effects while minimizing the induction of exhaustion markers and pro-tumoral macrophages.

### Immune cell fingerprinting of GC responses across cell-types revealed patterns of coregulation

Exploiting the structure-guided variance from the SGRMs, LPML generated a signaling fingerprint of GC action across immune and cancer cell subsets. A heatmap of the significant Pearson correlations, plotted as | r |, illustrates overall patterns of conserved and divergent coregulation from the repeated ligand perturbation (**Fig. 3A, fig. S5A**). As previously discussed, in addition to CD8^+^ T cells and CD4^+^ Th1 cells, we also differentiated three distinct CD4^+^ Th17 cell subsets. Th17 cells are known for their context-dependent roles in cancer and autoimmunity (**fig. S3**) (*26, 27*). Non-pathogenic Th17 cells (NP Th17), induced by TGFβ1 and IL-6 signals, promote tissue homeostasis, particularly in the gut, without causing autoimmunity. Pathogenic 1 (P1) and P2 Th17 subsets, each driven by IL-23 signaling, mainly differing in the requirement for TGFβ, show a gradient of increasing persistence and pro-inflammatory effects. Both subsets can induce disease in mice, with the severity of disease enhanced with P2 Th17 cells (*28*).

**Figure 3.**
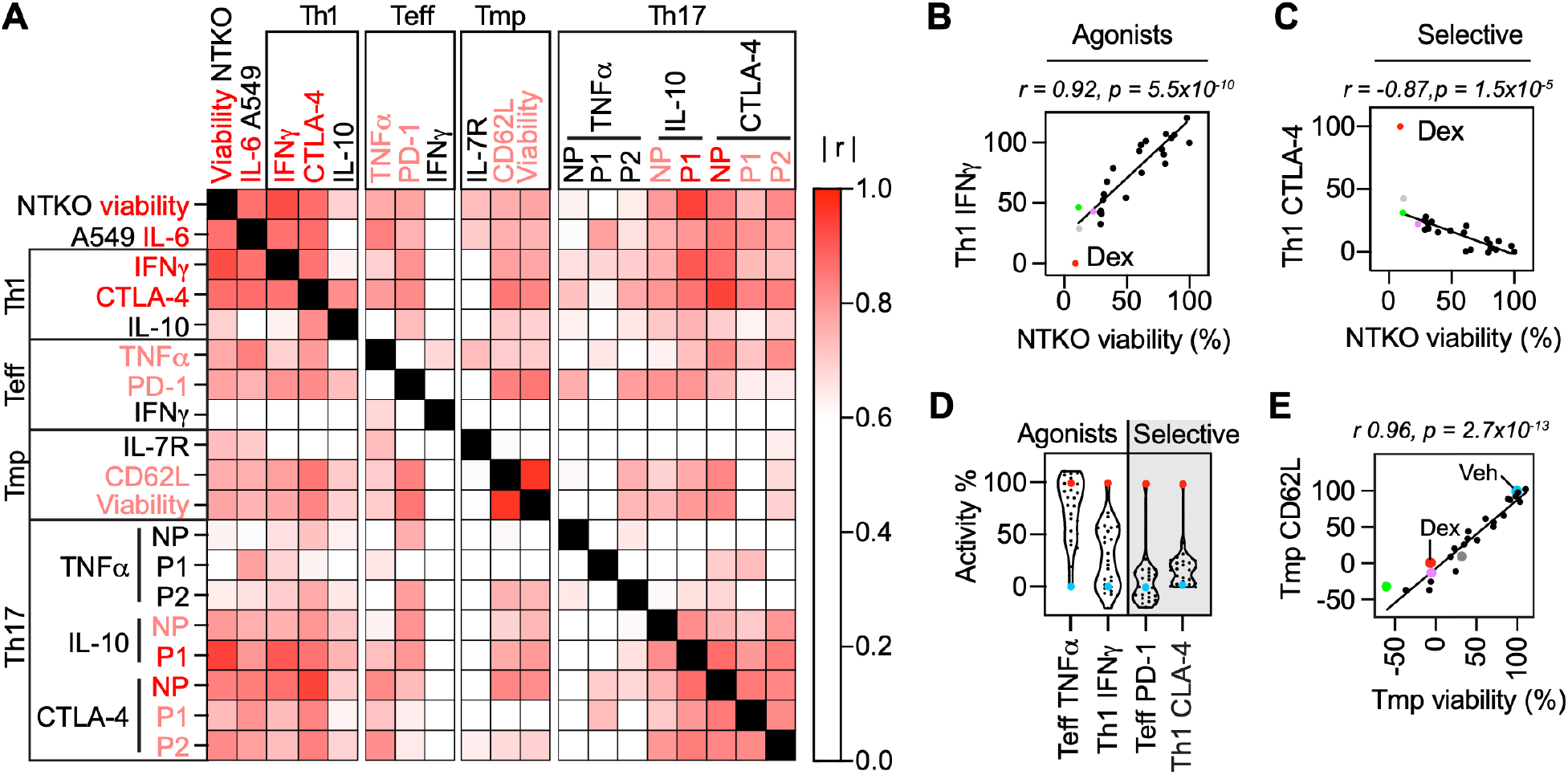
LPML reveals networks of GC-coordinated signaling in immune cells. **(A)** Heatmap of Pearson correlation matrix calculated from the SGRM effects in the indicated assays. Significance determined by non-zero slope test with Bonferroni corrected threshold of | r | > 0.66. Inclusion of assays was based on significant effects of Dex (*See Fig. 1–2*), and a Pearson correlation between biological replicates > 0.8, so that correlation between cell types or immune regulators was not limited by assay performance. See also **Fig. S5. (B–C)** LPML correlation plots Pearson correlation, r, calculated from SGRM activity excluding Dex and vehicle. Effects of SGRMs on NTKO cell viability and in the indicated assays in CD4+ Th1 cells. Each dot represents a separate compound, with indicated compounds colored as in **Fig.2.(D)** Violin plots displaying the distribution of SGRMs effects as % Dex. **(E)** LPML correlation plots Pearson correlation showing the effects of SGRMs on CD8+ Tmp cell viability or expression of CD62L.

SGRM effects on NTKO viability, A549 cell IL-6 secretion, Th1 cell INFγ production, and CTLA-4 expression were highly coregulated with each other and more widely across immune cell types and assays (**Fig 3A, Fig. 3B-C**). Effects on Teff PD-1 and TNFα were also significantly correlated with other cell types, including NTKO cell viability, reflecting broad GC coordination of inflammation and checkpoint function (**Fig. 3A, fig. 5B-C**). Effects on NTKO viability compared to IL-6 secretion in A549 lung cells or CD4^+^ and CD8^+^ T cell cytokines were similar, showing agonist activity from the SGRMs **(Fig. 1I, Fig. 3D, fig. S5B**). The high correlations were surprising for the checkpoint proteins, PD-1 and CTLA-4, where there was little SGRM increased protein expression (**Fig. 2B**). SGRM effects on PD-1 vs. NTKO viability showed a showed a tight clustering and significant correlation around the regression line despite the modest induction (**fig. S5C**). This shows the power of LPML to identify signaling patterns uniquely revealed by the structured guided variance of the ligands. Selective modulation in these contexts can be visualized in the skewing of the data distributions away from Dex (**Fig 3D, fig S6A–B**), separately from the correlation analysis. The coregulation of other GR-mediated functions with checkpoint proteins demonstrates sharing of a common signaling pathway, but also distinct differences driving the lack of checkpoint expression, suggesting Dex utilizes additional pathways for their expression or that the SGRMs drive repressive pathways. SGRM regulation of Tmp cell viability and CD62L protein expression were also widely coregulated with effects on other cell types (**Fig 3A, E**). Inhibition of viability was specific for Tmp and not Teff cells (**fig. S4B**), likely due to GCs effects on IL-2 signaling (*21*). Tmp viability was highly coupled to loss of CD62L, a selectin that drives lymph node homing (**Fig. 3E**), and surprisingly, not IL-7R (**Fig 3A**), a central mediator of T cell survival and persistence, particularly for memory T cell pools (*32*).

Comparing CD4^+^ T cell subsets differentiated and treated in parallel enabled LPML visualization of patterns across a gradient of inflammatory potential. IL-10 patterns in Th1 cells showed more restricted patterns of SGRM coregulation than the Th17 subsets (**Fig 3A-C, fig S6A–C**), despite Th17 cells expressing lower overall levels of IL-10 than the Th1 cells (*data not shown*). No IL-10 was detected in the P2 subset, reflecting the pathogenic potential of this cell subset. Effects on TNFα in the Th17 cells were not coregulated widely with other T cell subsets, while Th17 CTLA-4 showed tight regulation by the SGRMs across the cell types, and within the Th17 subsets, despite the limited activation compared to Dex (**Fig 3A, fig S6B –C**). Differences in GC regulation patterns for these critical immune modulatory proteins reflect the complexity of balancing secretion of pro- vs. anti-inflammatory mediators in T helper cells (*34*), illustrating how GCs tune homeostatic responses. Collectively, these data demonstrate the power of LPML to identify patterns in complex multi-dimensional data, providing computational attention to highly coregulated processes that are not apparent from individual compound profiling. GC action showed unexpected patterns of conservation, variance, and selective modulation across immune cell types and assays, demonstrating linkages between a specific allosteric signaling route and global coordination of cellular adaptations, such as immune cell phenotypes and metabolism. The LPML fingerprint confirms that while many GC responses are interconnected, specific allosteric inputs from the solvent channel can selectively decouple the core anti-inflammatory program from other immune modulatory effects.

### GCs coordinate checkpoint associated gene networks within CD8^+^ T effector cells

To uncover the molecular basis for these selective immunomodulatory effects, transcriptional profiling of CD8^+^ Teff cells provided the traditional benchmark of comparing the lead SGRM, SR38946, with Dex. The two compounds regulated a total of 773 genes (**Supplemental dataset 1)**: 652 genes by Dex and 538 genes by SR38946 (**Fig. 4A**), with substantial overlap between the target gene sets (**Fig. 4B**). SR38946 upregulated ∼70% of the number of Dex-induced genes but maintained a similar number of repressed genes (**Fig. 4A**). Gene set enrichment analysis (GSEA) of the molecular signatures database *Hallmark* gene set revealed that Dex- or SR38946-treated Teff cells upregulated *TNFα/NF-κB signaling* and *the inflammatory response*, and downregulated gene sets associated with cell proliferation such as *E2F targets*, as well as *Cholesterol homeostasis*. However, SR38946-treated cells limited or reduced upregulation of Dex-induced processes such as *apoptosis, IFNγ response*, and *IL-2/Stat5* and *IL-6/Stat3* signaling pathways (**Supplemental dataset 2**). Head-to-head comparison of the effects of Dex versus SR38946 on immunologic gene sets (*40*) revealed that compared to Dex, SR38946 reduced enrichment of several gene sets associated with CD8^+^ Teff cell function. For example, gene sets that are upregulated by CD8^+^ Teff cells in response to IFNγ stimulation were enriched by Dex but not SR38946 (**Fig. 4C, Supplemental dataset 2**). Also, gene sets that are downregulated by CD8^+^ Teff cells in response to IL-4, IL-7, or IL-18 stimulation were selectively enriched by Dex compared to SR38946 (**Fig. 4C, Supplemental dataset 2**). These results demonstrate that SR38946 circumvents adverse effects of GCs on CD8^+^ Teff cell function, particularly those linked to interferon signaling and cellular proliferation.

**Figure 4.**
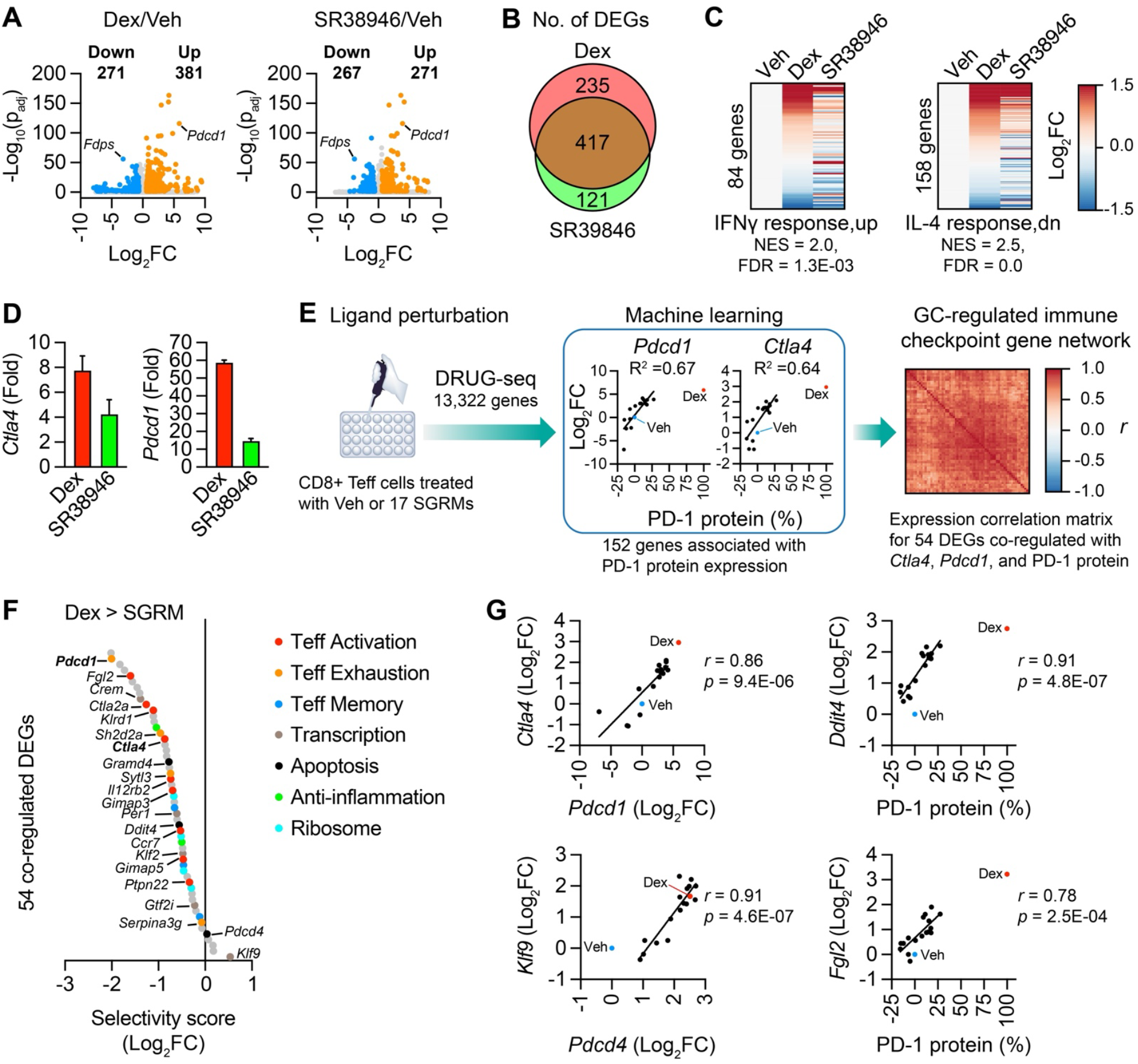
SGRMs selectively modulate a GC-regulated immune checkpoint-associated gene network in CD8+ Teff cells. **(A)** Volcano plots showing GC-dependent changes in the transcriptional profiles of CD8+ Teff cells treated with DMSO vehicle (Veh), 1 µM Dex, or 10 µM SR38946 (n = 3 independent replicates). The number of upregulated and downregulated genes (i.e., genes with a fold change ≥ 1.5-fold relative to vehicle and adjusted p-value (p_adj_) < 0.05) are also indicated. **(B)** Venn diagram showing the overlap between the differentially expressed genes (DEGs) in CD8+ Teff cells described in panel A. **(C)** Heatmaps comparing the expression profiles of the indicated CD8 T cell immunologic signature gene sets in cells described in panel A. Each row represents a distinct gene. The GSEA (Dex/SR38946) normal enrichment score (NES) and false-discovery rate (FDR) are also indicated for each gene set. **(D)** Upregulation of *Ctla4* and *Pdcd1* mRNA in CD8+ Teff cells described in panel A. Bars represent the average fold change relative to vehicle (mean + s.e.m). **(E)** LPML of the CD8+ Teff cell transcriptome revealed a co-regulated network of immune checkpoint-associated genes. Expression levels of *Ctla4, Pdcd1*, and 150 other genes were associated with PD-1 protein expression (simple linear regression, coefficient of determination, R^2^ ≥ 0.5). The heatmap shows a gene-by-gene expression correlation matrix for the 54 DEGs in the checkpoint-associated gene network (Pearson correlation coefficient, *r* ≥ 0.7 with PD-1, *Pdcd1*, and *Ctla4*, excluding Dex and vehicle). **(F)** Compared to Dex, SR38946 limits upregulation of the checkpoint-associated gene network in CD8+ Teff cells. Each datapoint represents a DEG, color-coded by their functional annotations. Selectivity score = SR38946 (Log_2_FC) – Dex (Log_2_FC). **(G)** Scatter plots of SGRM effects on the expression of select genes, showing expression selectivity for Dex or SR38946. Pearson correlation, r, calculated from SGRM activity, excluding Dex and vehicle.

Consistent with our immunophenotyping data, we noted that compared to Dex, SR38946 limited the upregulation of the immune checkpoint genes *Ctla4*, and *Pdcd1*, which encodes the PD-1 protein (**Fig. 4D**). LPML analysis of the CD8^+^ Teff transcriptome identified SGRM-responsive genes that are co-regulated with PD-1 protein expression, as well as *Pdcd1* and *Ctla4* mRNA levels, defining a GR-regulated immune checkpoint-associated network of ∼54 DEGs (**Fig 4D, Supplemental dataset 3**). Most of the genes in this network are involved in Teff cell activation and exhaustion, with a few genes involved Teff memory (**Fig. 4E**). Interestingly, five of these genes encode transcription factors i.e., *Crem, Klf2, Klf9, Per1*, and *Gtf2i* (**Fig. 4F**). As with the protein data, analysis of LPML correlation plots showed a fingerprint of gene associations. PD-1, *Pdcd1*, and *Ctla4* were highly co-regulated with reduced induction by the SGRMs (**Fig. 4D-F**). Few genes such as *Klf9* showed equal or greater activity with the SGRMs (**Fig. 4E-F**), similar to macrophages (**Fig 2F**).

We also examined other GC-regulated gene networks in CD8^+^ Teff cells. Fkbp5 is a primary effector of organismal GC responses, acting as a scaffold for various inflammatory mediators such as NF-κB and as a feedback inhibitor of GR function, but also mediate GC effects on metabolism including insulin resistance in skeletal muscle and muscle atrophy (*41, 42*). The *Fkbp5* module contains TCR inhibitory genes (*Ptpn1* and *Sla2*), and regulators of metabolic stress, survival, and tissue trafficking. The correlation matrix showed that core GC-regulated genes, *Fkbp5* and *Nr3c1* which encodes GR were highly co-regulated with PD-1 protein expression, but not with *Clta4* in the checkpoint-associated network, or *Il7r* in a distinct module (**Supplemental dataset J3**). This also applies to the correlated gene networks associated with each of these key GC driver genes (**Supplemental dataset J3**). The most highly co-regulated genes in the *Il7r* module also displayed high correlation with *Ctla4*, but not effects of the SGRMs on *Fkbp5* or *Nr3c1* expression. Functional annotation of these different gene sets showed that the SGRM effects on *Il7r* expression is co-regulated with genes that encode proteins involved in trafficking and quiescence, as well as chromatin regulatory factors. Together, these results demonstrate that LPML enables identification of gene regulatory networks based on real perturbation studies, with the structured variance of the ligands driving the identification of coregulation.

### Molecular dynamics simulations revealed leveraged allostery from the solvent channel

**TBD**

## Discussion

This structure-based design strategy built upon the synthesis of novel steroidal scaffolds described by Kamenecka et al. (*43*) and the subsequent biological characterization of the prototype modulator, SR11466(*9*). This previous work demonstrated that engaging the solvent channel could mechanistically dissociate skeletal muscle atrophy from anti-inflammatory activity, validating that the solvent channel acts as a druggable allosteric hub. Here, we describe expansion upon the chemical series to focus on the GR solvent channel in conjunction with our LPML model to identify a series of compounds to maintain optimal anti-inflammatory Dex-like effects while minimizing some of the negative effects observed on immune responses previously observed with GC treatment.

Development of a high-throughput T cell profiling platform enabled us to separate anti-cancer from anti-inflammatory effects and T cell exhaustion markers. The library of solvent channel-binding SGRMs included compounds that retained the ability to induce apoptosis in multiple myeloma cells and to suppress inflammatory cytokine production (IL-6, TNFα) with efficacy comparable to that of Dex. These same compounds attenuated the induction of expression of T cell exhaustion markers, specifically PD-1 on CD8^+^ effector T cells and CTLA-4 on CD4^+^ helper T cells. This divergence has challenged the long-standing dogma that GR-mediated immunosuppression is a uniform phenotype for most corticosteroids (*44*). Using our LPML framework, we showed that cancer cell death and cytokine suppression programs are separable from checkpoint induction, suggesting that immunosuppressive effects may arise from specific allosteric states that can be selectively avoided. This aligns with recent findings that glucocorticoid sensitivity is not uniform but relies on cell-type specific thresholds and cofactor availability in macrophages versus stromal cells (*45*).

As an attention mechanism, LPML is not sufficient to assign causality, but rather enriches in potential mediators that we identified in a small-scale gene perturbation study. By studying responses across a series of ligands, LPML amplifies signals from coregulated pathways over transcriptional noise. This powerful correlation, however, does not prove that a given gene is a functional driver. For instance, a gene’s expression might be tightly coupled to checkpoint induction because both are downstream targets of a shared master regulator, not because one directly influences the other. The true utility of LPML is in converting an intractable ‘big data’ problem into a testable set of hypotheses. It provides a high-confidence shortlist of candidate genes as causal nodes within the network.

GCs present a paradox in oncology. The direct tumor-killing effects of dexamethasone is countered by it induction of T cell exhaustion, which can antagonize the efficacy of checkpoint inhibitors such as anti-PD-1/PD-L1 therapies (*47, 48*). A ligand that decouples these activities, killing the tumor without disarming the immune system, could present an ideal therapeutic candidate. Beyond oncology, this selectivity profile offers a roadmap for safer maintenance therapies in chronic autoimmune diseases, where the goal is to control specific inflammatory pathways without increasing patients’ vulnerability to opportunistic infections. Here, we establish the GR solvent channel as an allosteric control node, validating it as a target for design of immune-selective SGRMs that decouple anti-tumor efficacy from T cell exhaustion, and the GR signaling code was shown to be pliable and context-dependent. By achieving selectivity at the level of receptor conformation rather than rapid metabolism, this approach could enable safer systemic therapies.

## Acknowledgements

This work was supported by the grants to K.W.N. – NIH R01 GM146385; I.R. - NIH R01DK099087, NIH R01AI148129 and The Hospital for Special Surgery David Rosensweig Genomics Center; J.D.L – Blood Cancer United Specialized Center Research Grant 7021-20, Myeloma Solutions Fund IMMPACT award; L.A.S. - NIH R01 GM146385, R01DK135300, R01DK136298; T.M.K. - NIH R01 GM146385.

## Author Contributions

K.W.N. conceived the project. K.W.N, T.M.K., R.R.K, J.C.N., L.A.S., I.R., J.D.L., L.S., T.P.S., M.J., A.S. designed research. R.R.K., C.K.M., J.C.N., H.Z., L.B., B.S., W.J.S., C.M., J.S., E.B., A.S., A.R.T., M.K. performed research. K.W.N., R.R.K., J.C.N., C.K.M., T.T.V, M.E.P., T.M.K., W.J.S., I.R., C.M., T.P.S., L.A.S. analyzed data. K.W.N., L.A.S, J.C.N., C.K.M., and R.R.K. wrote the manuscript with edits and input from all others.

## Methods

### Mice

For all *in vitro* experiments, both male and female C57BL/6 mice were either purchased from the Jackson Laboratory (Bar Harbor, ME) or Charles River Labs (Charleston, SC) and/or bred at the Wertheim UF Scripps Biomedical Research Institute. Mice, five to ten weeks old, were sacrificed during the light cycle. All mice were maintained under specific pathogen free conditions at controlled temperature (22–23 C), humidity 60%, and 12h:12h light:dark cycles. Mice had access to regular chow (Harlan 2920X) and water, *ad libitum*. To generate mice with a T cell-specific deletion of the GR, GR floxed mice (Jackson labs, Bar Harbor, ME; B6.Cg-Nr3c1tm1.1Jda/J) were crossed with dLCK-Cre transgenic mice (Jackson lab, Bar Harbor, ME; B6.Cg-Tg(Lck-icre)3779Nik/J). All studies conform to the guidelines of and were approved by the Wertheim UF Scripps Biomedical Research Institutional Animal Care and Use Committee (IACUC) or in compliance with the Weill Cornell Institutional Animal Care and Use Committee (IACUC).

### Chemical Synthesis

Compounds, including SR11466, were synthesized as previously described (*9, 43*).

### Luciferase Co-transfection Reporter Assays

HEK293-T cells obtained from American Type Culture Collection (ATCC) were cultured in Dulbecco’s minimum essential medium (DMEM) (Cellgro by Mediatech, Inc. Manassas, VA) supplemented with 10% fetal bovine serum (FBS) (Hyclone by Thermo Scientific, South Logan, UT), and 1% non-essential amino acids (Cellgro), Penicillin-Streptomycin-Neomycin antibiotic mixture and Glutamax (Gibco by Invitrogen Corp. Carlsbad, CA), were maintained at 37°C and 5% CO2. HEK-293T cells were seeded in a 10-cm dish containing 10 ml of phenol red-free DMEM (catalog no. 31053028; Thermo Fisher Scientific) containing 10% charcoal-stripped FCS (catalog no. A3382101; Thermo Fisher Scientific). The next day, cells were cotransfected with 10 µg of MMTV-luciferase reporter plasmid and 5 µg of steroid receptor plasmid, using 45 µl of TransIT-LT1 reagent (catalog no. MIR 2300; Mirus Bio). After 24 h, cells were transferred to a 384-well plate (catalog no. 781080; Greiner Bio-One). The next day, the test compounds were added using a Biomek NXP 100-nl pintool (Beckman Coulter). Luciferase activity was measured 24 h later using the britelite plus Reporter Gene Assay System (catalog no. 6066761; PerkinElmer).

### CD8^+^ Cells

Naïve CD8^+^ T cells were isolated from spleens and lymph nodes of mice using the EasySep Mouse Naïve CD8+ T cell isolation kit (STEMCELL Technologies, Canada, 19858A) following the manufacturer’s instructions after removing red blood cells using Lympholyte-M (Cedarlane Laboratories). Cells were cultured in T-cell medium (Dulbecco’s modified Eagle’s medium supplemented with 10% FBS, non-essential amino acids, 10 mM HEPES, 2 nM L-glutamine, sodium pyruvate, arginine, aspartate, folic acid, 50 µM 2-mercaptoethanol, MEM vitamin solution) at 37°C in 10% CO2 in a volume of 200 μL in 96-well U-bottomed plates. After isolation, cells were plated at a density of 2×105 cells per well and activated with anti-CD3 (clone 2C11, 1 μg/mL) and anti-CD28 (clone 37.51, 1 μg/mL) by precoating plates with 50 μg/mL goat anti-hamster IgG (MP 56984). After 48 hours, the activated cells were directly resuspended, counted, and re-cultured at a concentration of 1×105 cells per well by diluting the cells into fresh media containing 100 U/mL recombinant human IL-2 (NIH) to generate effector-like cells; cells were re-cultured daily. On day 5, cells were treated with 1 μM of compound or DMSO (vehicle), and on day 6, they were restimulated with 50 ng/mL phorbol-12-myristate-13-acetate (Sigma) and 1 μg/mL ionomycin (Sigma) for 2 h with the addition of GolgiStop (BD Bioscience) for an additional 2 h before staining. Following staining for viability, cells were fixed and permeabilized using the Foxp3 staining kit (eBioscience) with anti-IFNγ and anti-TNFα. Staining was performed using FACS buffer (0.5% BSA, 2mM EDTA in PBS). All flow cytometry data were analyzed using a BD Symphony instrument (BD Biosciences) and FlowJo 10.10.0 software. Data are presented as means ± SEM, and statistical significance was determined using independent two-tailed Student’s t-tests in GraphPad Prism 10.

### CD4^+^T Cells

Naïve CD4^+^ T cells from spleen and lymph nodes of mice were purified after removing the red blood cells using Lympholyte-M solution (Cedarlane Laboratories). Cells were enriched for naïve CD4+ T cells using the mouse naïve CD4^+^ T Cell Isolation Kit (STEMCELL Technologies, Canada) according to the manufacturer’s instruction. Cells were cultured in IMDM medium (Invitrogen) with 10% FBS, 100 IU/mL penicillin, 100 μg/mL streptomycin, 50 μM β-mercaptoethanol, and 2 mM L-glutamine. All cultures were performed in a volume of 200 μL in 96-well U-bottomed plates. The conditions for the different T helper cell subsets were: For Th1 conditions: 5 μg/mL anti-IL-4, 20 ng/ml IL-12 (R&D Systems), and 10 ng/mL IFNγ (R&D Systems); For Th17 conditions: 5 μg/mL anti-IFNγ, 5 μg/ml anti-IL-4 (R&D Systems), and 1.5 ng/mL TGFβ (R&D Systems) and 30 ng/mL IL-6 (R&D Systems). Other cytokines used for Th17 conditions: IL-1β (10 ng/mL, R&D Systems) and IL-23 (20 ng/mL, R&D Systems). 1×10^6^ cells/mL of naïve CD4+ T cells were activated with anti-CD3 (clone 2C11, 0.5 μg/ml) and anti-CD28 (clone 37.51; 1 μg/ml) by precoating plates with 50 μg/mL goat anti-hamster IgG. After 48 h, cells were removed from the TCR signal and recultured at a concentration of 1×10^6^ cells/mL. Th1 cells were cultured with addition of recombinant IL-2 (10U/mL; NCI, Biological Resources Branch) at 48 h and 72 h. Four days after activation, all cells were restimulated with 50 ng/mL phorbol-12-myristate-13-acetate (PMA) (Sigma) and 1 μg/mL ionomycin (Sigma) for 2 h with the addition of GolgiStop (BD Bioscience) for an additional 2 h before intracellular staining.

### T cell Drug Treatment and Flow Cytometry

The compounds were added to the T cells using a Biomek NXP 200-nl pintool (Beckman Coulter), treating the cells with 200 nL of stocks into 200 μL of media for a final concentration of 10 μM for the CD8^+^ T cells and 1 μM for the CD4^+^ T cells based on dose-response curves with dexamethasone and representative compounds (*data not shown*). For CD8^+^ T cells, drugs were added on day 5 and for CD4^+^ T cells on day 3 of differentiation, both for a duration of 16-24 h. Cells were washed with PBS and surface stained with fluorescence-conjugated antibodies for 30 minutes at 4C in the dark, washed with PBS, then resuspended in FACS buffer (0.5% BSA, 2 mM EDTA in PBS). For intracellular cytokine staining, cells were restimulated with 50 ng/mL phorbol-12- myristate-13-acetate (PMA) (Sigma) and 1 μg/ml ionomycin (Sigma) for 2 h with the addition of GolgiStop (BD Bioscience) for an additional 2 h. Cells were then surface stained using procedures outlined above, fixed and permeabilized using the Foxp3 staining kit (eBioscience) at 4C in the dark for 45 minutes, followed by intracellular cytokine staining at 4C in the dark for 30 minutes.

### Macrophage Polarization

Bone marrow cells were isolated from 6-weeks-old C57BL/6J mice and seeded at a density of 6×10^5^ cells/mL in DMEM (10% FBS, and 1% PenStrep, Gibco) with 10 ng/ml recombinant mouse colony stimulating factor 1 (mCSF-1) (Gibco, PMC2044) in a 12-well flat bottom plate (Corning, 351143). After 4 days, DMEM containing 10 ng/mL MCSF-1 was replenished. Macrophages were then treated with 10μM of vehicle, Dex, or GR compounds for 48 h. Then, the differentiated macrophages were removed from the culture plate by adding 500 μL Accutase™ (Stemcell Technologies, 07920) to each well and incubating at 37C for 10 minutes. Macrophages were washed once with DMEM and resuspended in 100 μL of PBS. The cells were then stained with fixable viability stain FVS520 for 20 minutes at RT, followed by the immunostaining (1 hour at 4C) with for extracellular proteins: CD11c, F4/80, and CD11b. Cells where then washed with PBS and resuspended in 200 μL of Fixation/Permeabilization working solution (Invitrogen, 00-5123-43, 00-5223-56) and incubated at 4C for up to 16 hours. Next, they were washed with Permeabilization Buffer 1X (Invitrogen, 00-8333-56) and stained for intracellular CD206 for 1 hour at 4C, and washed once with Permeabilization Buffer 1X (Invitrogen, 00-8333-56), once with PBS and resuspended in 200 μL of PBS. Samples were analyzed on a BD LSR II cytometer (BD Biosciences). For BMDCs profiled for qRT-PCR, BMDMs were prepared as previously described (Deochand et al, *Nature Comms* 2024). Briefly, bone marrow was flushed from the femurs and tibiae of mice and differentiated for 5 days in the DMEM (1 g/l glucose) supplemented with 20% bovine serum and 10% L-cell conditioned media. Adherent cells were scraped, seeded into 12-well plates and polarized for 24 h with 10 ng/ml IL4, with 100 nM Dex or indicated ligands (10μM) added for the last 8 h.

### RNA isolation and qRT-qPCR

Total RNA from macrophages treated as described in Figure legends was isolated using Qiagen RNAeasy Plus Mini kit. RNA samples were subjected to random-primed cDNA synthesis and gene expression was analyzed by qPCR with Maxima Sybr Green/ROX/2x master mix (Thermo Scientific™) on StepOne Plus real time PCR system (Applied Biosystems) using standard protocols, and data was analyzed using ΔΔCt method with b-actin transcript as housekeeping control. Primers used are as follows: *mActb* Fw: AGGTGTGCACTTTTATTGGTCTCAA; *mActb* Rv: TGTATGAAGGCTTTGGTCTCCCT; *mKlf9* Fw: CCTTGGTGTTAGCTGAGATTG; *mKlf9* Rv: GAACAGGGAGGGAGAGAGAAA; *mArg1* Fw: ACCACACTGACTCTTCCATTCTT; *mArg1* Rv: TTGATGTCCCTAATGACAGCTCC; *mCd163* Fw: AAACCCTAAACGATTGGCTGC; *mCd163* Rv: ACTCTTCCTAAGCATCGGTGG.

### Flow Cytometry

Flow cytometric analysis was performed on a BD Symphony (BD Biosciences) instrument and analyzed using FlowJo software (TreeStar). All antibodies used in experiments are described in Supplementary Table X. Gating and subpopulation analysis was performed using FlowJo software (BD Biosciences). Reported percentages are given as the percentage of a parental gate. Singlets discrimination was based on FSC and SSC scatters, and dead cells were excluded by viability staining. This research project was supported in part by the Herbert Wertheim UF Scripps Institute for Biomedical Innovation and Technology Flow Cytometry Core Facility at the University of Florida (RRID:SCR_027278).

### IL-6 Secretion

A549 cells (ATCC) were cultured in high-glucose DMEM supplemented with glutamine and 10% FCS. For steroid-free culture conditions, cells were grown in charcoal:dextran-stripped FCS (catalog no. 100-119; Gemini Bio). IL-1β (R&D Systems) was dissolved in PBS plus 0.1% BSA (Sigma-Aldrich). A549 cells were plated in DMEM containing charcoal-stripped serum in 384-well plates. The next day, cells were treated for 6 h with compounds at 1 μM and then treated with 1 ng/ml1 of IL-1β for 6 h. Supernatants were collected, and IL-6 was assayed with AlphaLISA (PerkinElmer) according to the manufacturer’s instructions.

### C2C12 Myoblast Proliferation Assays

C2C12 cells were seeded in 384-well microplates containing 25 µl per well of phenol red free DMEM supplemented with 10% charcoal-stripped FBS. The cells were treated with compounds (dose curves) using a 100-nL pintool Biomeck NXP workstation (Beckman Coulter Inc.). Five days later, 25 µl per well of CellTiter-Glo reagent (Promega) was added, and luminescence was measured to assess cell viability. Data was normalized using cell numbers estimated from growth curves.

### DRUGseq

For transcriptional profiling T cells were prepared for the MERCURIUS™ DRUG-seq service (*49*) (Alithea Genomics, Frederick, MD). 45,000 cells were plated per well. T cells were treated with 1 µM of each test compound or 10 µM of dexamethasone (Dex). The T cells were cultured in 24-well plates, and 24-hour treatments were started on day 5 of culture in a final concentration of 0.1% DMSO. Following treatment, cells were washed, and plates containing frozen cell pellets were shipped to Alithea Genomics for processing. The service utilizes a highly multiplexed, extraction-free protocol where library preparation is performed directly from cell pellets. Triplicate wells for each condition were then pooled prior to library preparation, except samples treated with Vehicle (DMSO), Dex, SR38946, SR38951, and SR42364, which were processed as individual replicates. Alithea Genomics performed library quality control, sequencing, and initial data processing, providing raw sequencing data (fastq files) and gene count matrices. Raw and processed RNA-seq datasets will be deposited to the Gene Expression Omnibus (GEO), with accession number XXX, in the future. Downstream differential expression analysis was performed using DESeq2 (v1.50.2). Gene set enrichment analysis (GSEA) was performed using the *GSEA* software v4.3.3(*50, 51*). Gene sets analyzed, including the mouse orthology-mapped “hallmark” gene sets, and the immunologic signature collection (M7) were downloaded from the molecular signatures database (https://www.gsea-msigdb.org/gsea/msigdb/index.jsp). Gene sets with large gene sets with >500 genes and small gene sets with <15 genes were excluded from the analysis.

### Machine Learning

GC-dependent effects on PD-1 protein expression in CD8^+^ Teff cells was compared to log2-transformed fold changes in expression of 13,322 genes quantified in the CD8^+^ Teff cell transcriptome by DRUG-seq, through simple linear regression using the basic R function “lm” to fit linear models for each gene. The coefficient of determination, R^2^ reflects the strength of the linear relationship between the effects of the compounds on a gene and their effects on PD-1 protein expression. A threshold of R^2^ ≥ 0.5 was used to identify genes that are associated with PD-1 protein expression. This method was also to identify genes that were co-regulated with *Ctla4, Pdcd1* and other genes. Functional annotation of genes in the immune checkpoint-associated network was performed using the DAVID webserver (*52*) and previously published data sets (*53, 54*).

### NTKO/TKO Proliferation (Claudia in Spicer lab)

NTKO and TKO cells in suspension were cultured in Advanced RPMI 1640 Medium supplemented with 10% FBS, GlutaMAX, and an antibiotic/antimycotic solution. The cells were maintained at a seeding density of 2.5 x 10^5^ cells/mL and passaged two to three times per week. For the cell viability assay, 30 µL of cells in their maintenance media were dispensed into either 384- or 1536-well plates, followed by the addition of 100 nL of compound via pin transfer. After a 72-hour incubation at 37°C, the cells were lysed by adding 30 µL of CellTiter-Glo reagent and incubating for 10 minutes at room temperature. The plates were then centrifuged at 1200 rpm for 2-3 minutes, and luminescence was recorded on a PheraSTAR plate reader.

### Docking

### Molecular Dynamics Simulation

### Quantification and Statistical Analyses

All statistical analyses were performed using Prism v10 (GraphPad Software) unless otherwise stated. Data are presented as mean ± SEM. A two-tailed unpaired Student’s t-test was used for comparisons between two groups.

For comparisons of more than two groups, a one-way ANOVA was used, followed by Dunnett’s multiple comparisons test. Specific p-values are indicated in the figures, with significance denoted as *p < 0.033, **p < 0.022, and ***p < 0.001. To analyze signaling relationships, Pearson correlation coefficients (r) were calculated for the maximal efficacy of each SGRM across the functional assays and significance determined by the non-zero slope test from simple linear regression. Dose-response curves were fitted using three-variable non-linear regression. For the Pearson correlation matrix in fig. 3A, significance was determined by the Bonferonni corrected p value = 0.05/ 200 tests, producing an r threshold of 0.66 for significance.

---

**Table S1.** Luciferase data for all ligands evaluated.

| Compound | Emax (%) |  | EC50 (nM) |  | IC50 (nM) |  |
| --- | --- | --- | --- | --- | --- | --- |
|  | GR | PR | GR | PR | GR | PR |
| Dexamethasone | 98.11 | 43.18 | 0.05 | 51.86 |  |  |
| SR38944 |  |  |  |  |  |  |
| SR38945 | 101.2 |  | 35.18 |  |  | 366.60 |
| SR38946 | 95.91 | 52.06 | 0.23 | 1.31 |  |  |
| SR38947 | 94.22 | 25.86 | 2.58 | 14.10 |  |  |
| SR38950 | 85.81 | 44.83 | 0.43 | 0.57 |  |  |
| SR38951 | 77.35 | 61.09 | 5.08 | 0.92 |  |  |
| SR38952 | 90.46 | 36.22 | 46.82 | 560.00 |  | 62.26 |
| SR38953 | 91.68 | 42.9 | 27.22 | 4.60 |  |  |
| SR38954 | 94.99 |  | 2.11 | 56.60 |  | 9.14 |
| SR38955 |  |  |  |  |  | 206.40 |
| SR38956 | 51 |  | 461.60 |  |  | 48.65 |
| SR38957 | 89.66 | 61.01 | 3.04 | 1.90 |  |  |
| SR38964 |  |  |  |  | 36.78 | 0.0046 |
| SR41244 | 93.69 | 42.71 | 1.33 | 3.43 |  |  |
| SR42164 |  |  |  |  | 36.33 | 0.16 |
| SR42184 | 22.27 |  |  |  |  |  |
| SR42204 | 65 |  | 0.08 | 1.10 |  | 1.32 |
| SR42224 | 27.35 | 56.32 | 0.32 | 1.08 | 29.16 |  |
| SR42244 |  |  |  |  |  |  |
| SR42264 | 44.97 | 40.75 | 1.73 | 1.75 |  | 22.54 |
| SR42284 | 49.44 | 39.85 | 0.19 | 41.63 |  |  |
| SR42304 | 66.8 | 42.51 | 0.12 | 2.89 |  | 6.00 |
| SR42324 |  |  | 7.43 | 8.37 |  | 28.02 |
| SR42344 | 81.79 | 60.62 | 1.08 | 3.31 |  |  |
| SR42444 |  |  |  |  |  | 1084.00 |
| SR42464 | 93.32 |  | 0.28 | 9 |  | 214.50 |
| SR42484 | 78.43 |  | 768.70 | 460.00 |  |  |
| SR42364 | 102.1 | 42.93 | 0.11 | 0.25 |  |  |
| SR42384 | 92.36 | 51.05 | 0.31 | 0.92 |  |  |
| SR42404 | 103.6 | 40.21 | 1.40 | 1.30 |  |  |
| SR42424 |  |  |  |  | 17.64 | 0.13 |

## Supplementary Figures

**Figure S1.**
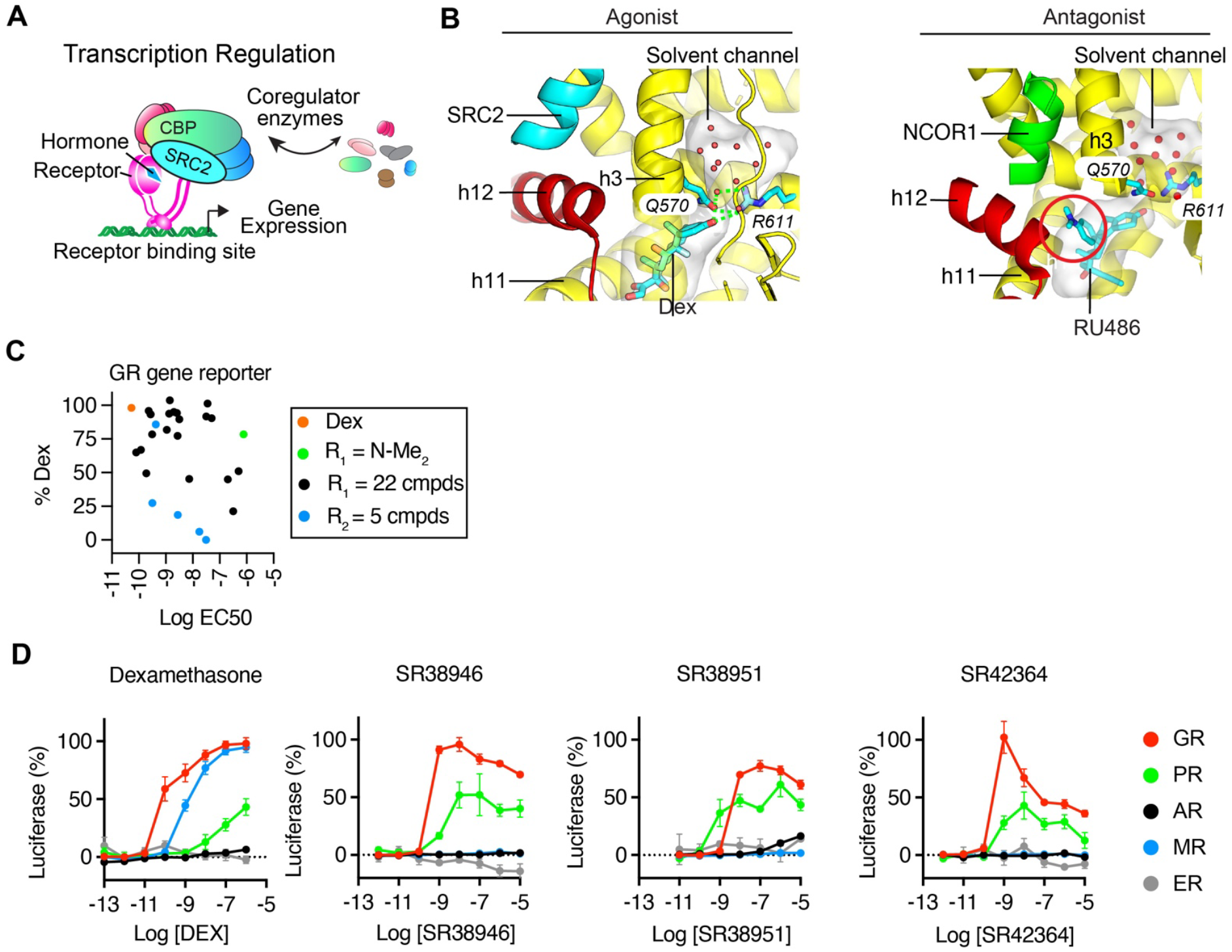
Structure based design and compound characterization. **(A)** Transcriptional regulation by nuclear receptors is through ligand mediated interaction with DNA and the dynamic recruitment of transcriptional coregulators that modulate gene expression. **(B)** Helix 12 controls the shape of the coregulator binding site and recruitment of different coregulators. Structures of GR bound to Dex or the GR/PR antagonist RU486, with the ligands shown as cyan sticks. The bound peptides from the SRC2/Grip1 coactivator or NCOR1 corepressor are shown. Waters are shown as red spheres. The ligand binding cavity and solvent channel are indicated with a transparent surface. The red circle indicates where the antagonist side chain of RU486 shifts the position of helix 12 (h12) to change the shape of the surface. Ligands can also regulate the dynamics of helix 12 through shifts in helices 3 and 11 (PMIDs: 9875847, 9808622). **(C)** Luciferase gene reporter activity showing the EC_50_ and maximal efficacy for GR activity. GR expression plasmid and MMTV-Luciferase were transfected into 293T cels and treated with the indicated ligands. Mean = 6 from 2 biological replicates. **(D)** Luciferase reporter assays comparing the dose-response profiles of the indicated GCs across the steroid receptors. (GR; Glucocorticoid receptor; PR, Progesterone receptor; AR, Androgen receptor; MR, mineralocorticoid receptor; ER, Estrogen receptor). N=6/assay.

**Figure S2.**
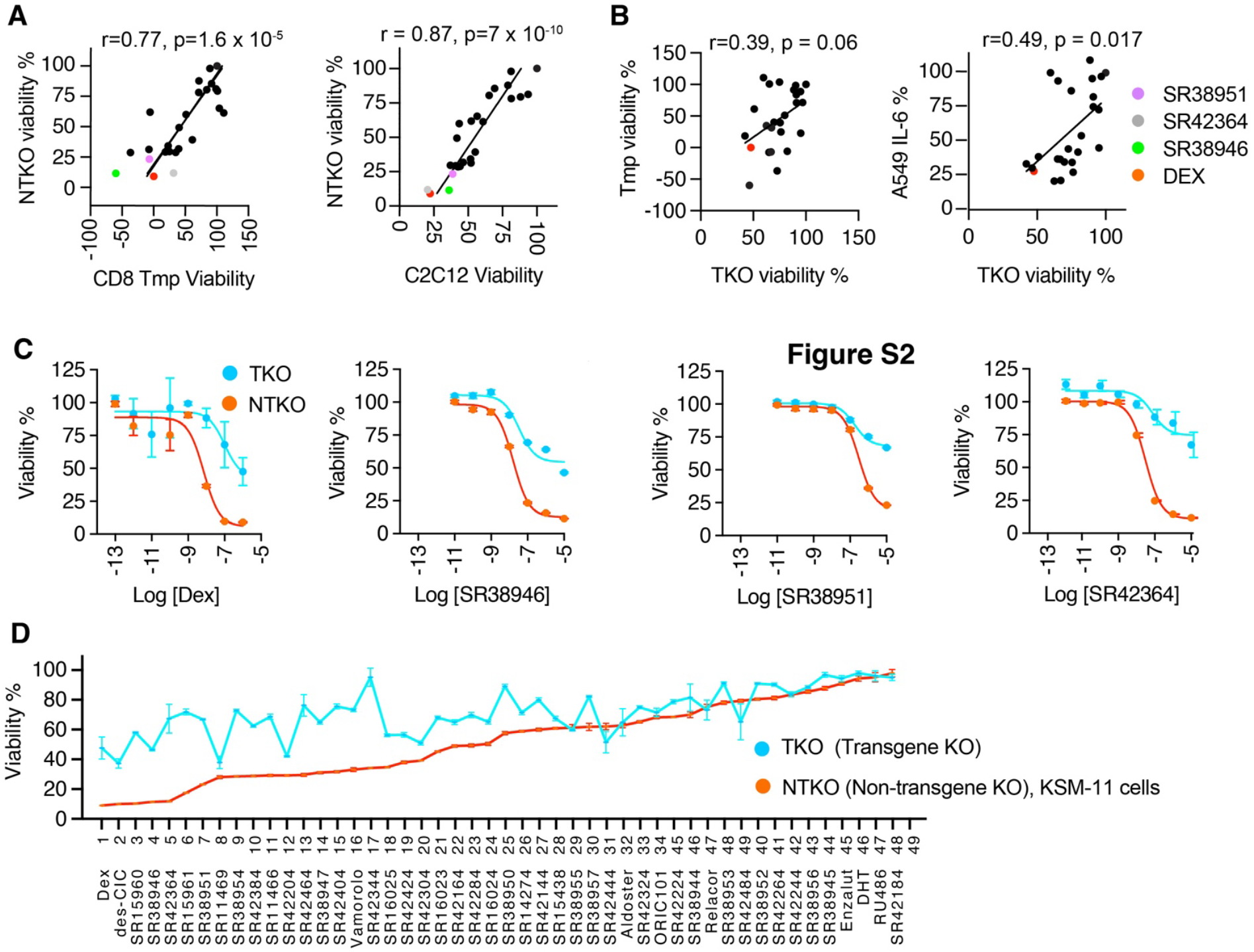
Scatterplots of effects of the R_1_-modified GCs on cell viability. **(A)** Inhibition of CD8 Tmp viability was determined by flow cytometry. C2C12 myoblasts were treated with dose curves of the GCs for 5 days and viability assessed with CellTiterGlo (Promega). Each dot represents the mean of effects of the compound at 10 μM (or 1 μM Dex) from two biological replicates of 2-4 wells each. Note that Dex has very different effects on myoblasts in differentiation media, promoting nuclei fusion and producing bigger myotubes, before later induction of atrophy (PMID: 33510451) **(B)** Scatterplots of viability of Tmp or C2C12 cells compared to TKO steroid resistant cells. **(C)** The NTKO or TKO cells were treated with dose curves of the compounds for 72 h and viability assessed with CellTiterGlo (Promega). N = 8, mean ± SEM. **(D)** Maximal effects of compounds as assayed in (C).

**Figure S3.**
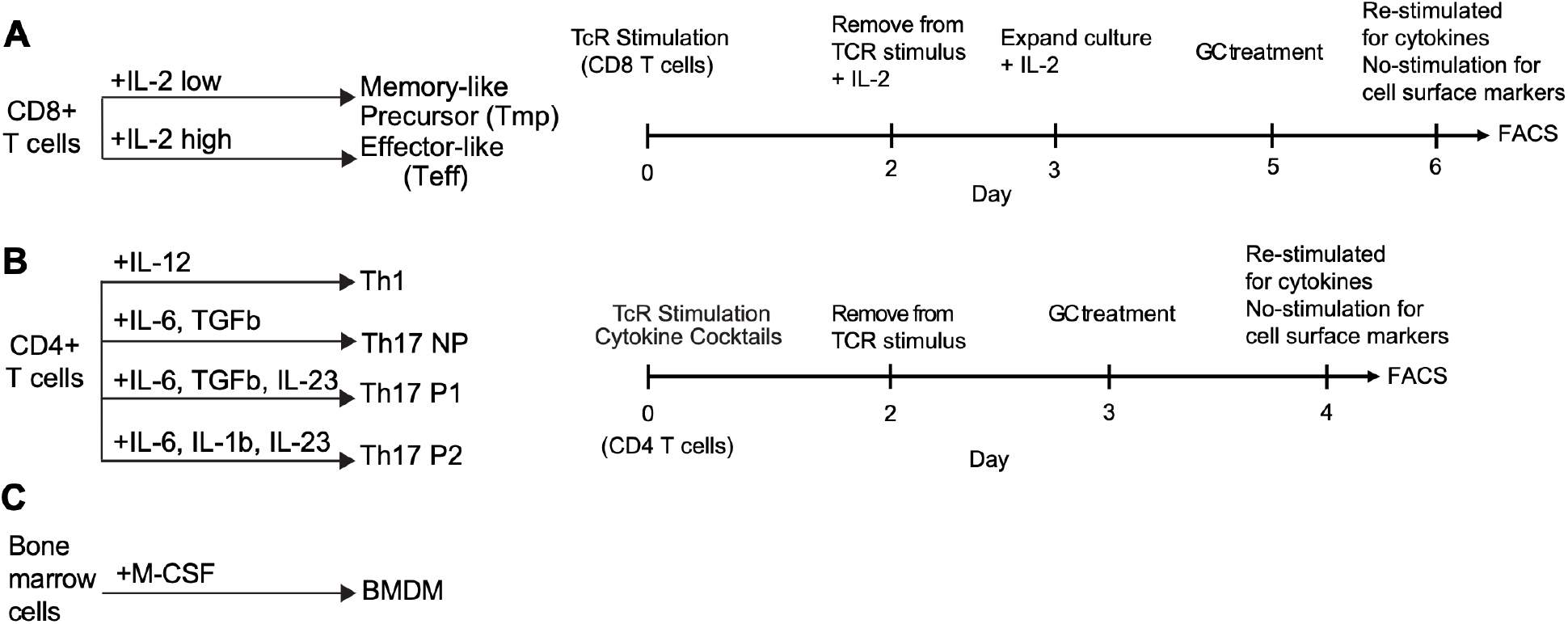
Conditions used for the *in vitro* differentiation of mouse T cell subsets and bone marrow-derived macrophages (BMDMs). (**A)** CD8+ T cell differentiation conditions. **(B)** CD4+ T cell differentiation conditions. **(C)** Differentiation conditions for bone-marrow derived dendritic cells (BMDCs).

**Figure S4.**
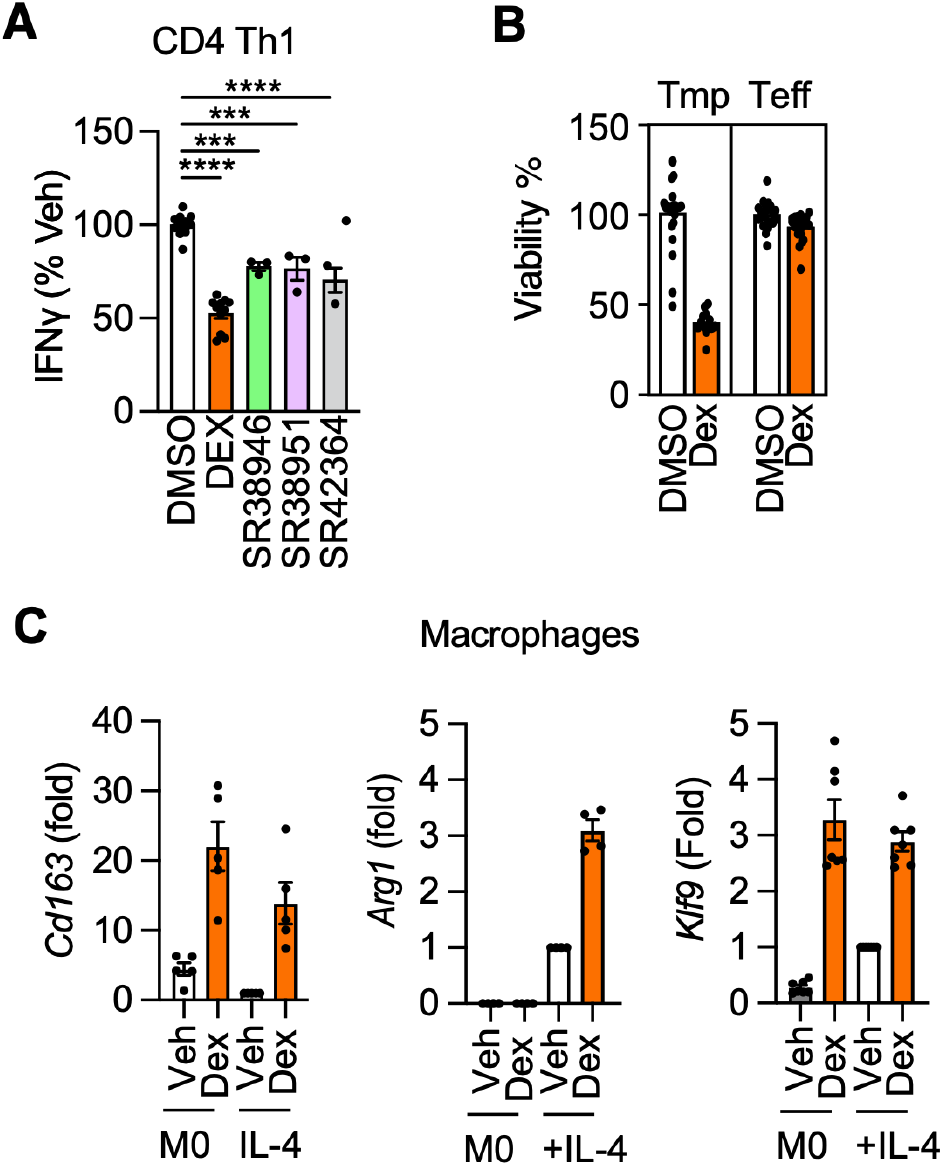
Effects of Dex or SGRMs on immune cell subsets. **(A)** IFNγ expression in CD4+ Th1 cells treated with the indicated GCs. Graph represents data normalized to vehicle or Dex treatments. n = 4-16, Mean + SEM. **(B)** Percent viability of CD8+ Tmp vs Teff cells normalized to vehicle vs Dex treatment. n = 4-16, Mean + SEM. **(C)** Expression of *Cd163, Arg1*, and *Klf9* in BMDMs differentiated in the absence or presence of IL-4 in addition to vehicle or Dex treatment for the last 8h. n = 4-7, Mean + SEM. Significance determined by one-way ANOVA with Dunnett’s multiple comparisons test (* p < 0.033, ** p < 0.022, *** p < 0.001).

**Figure S5.**
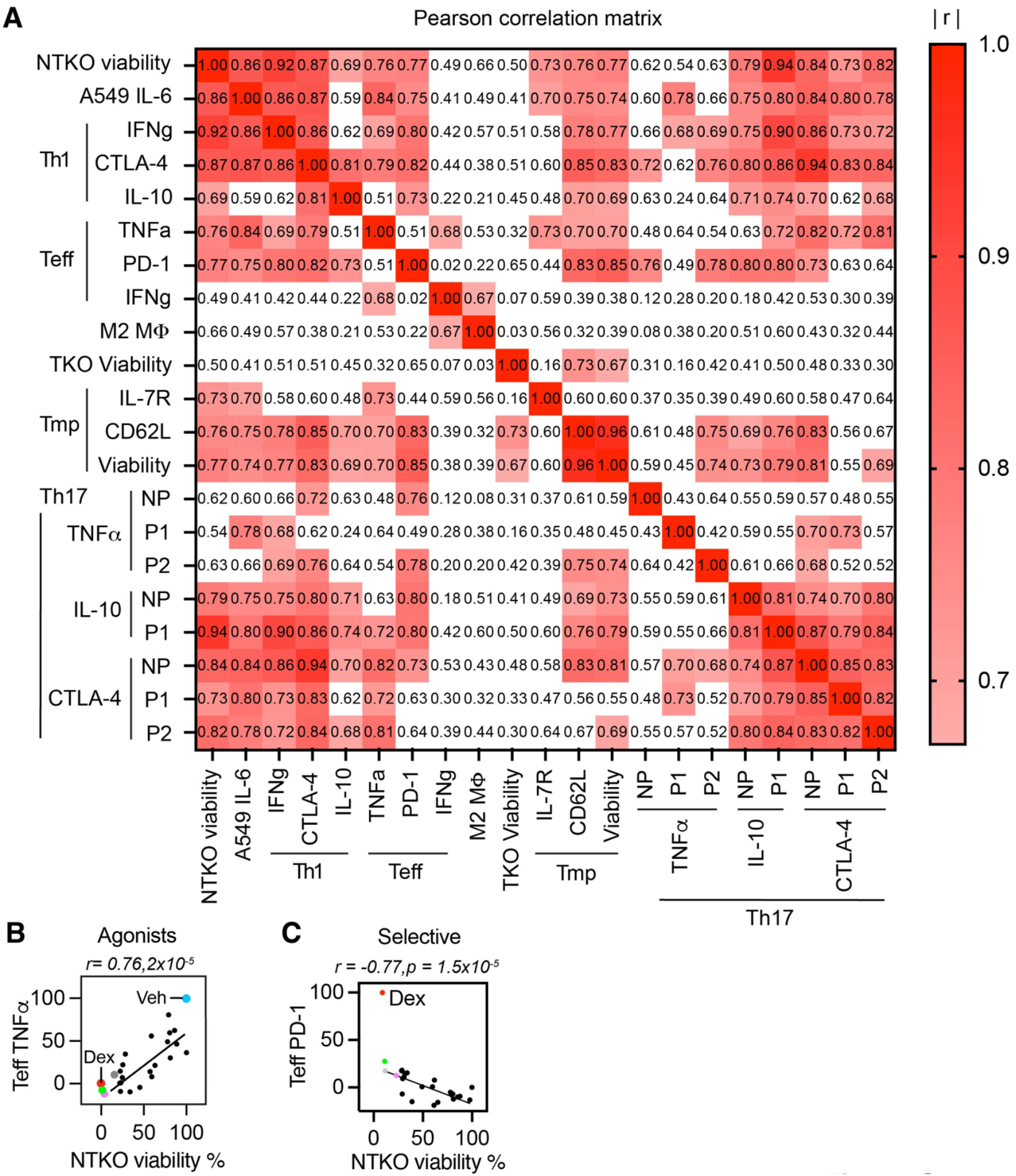
Heatmap of Pearson correlation matrix. **(A)** Heatmap of Pearson correlation matrix calculated from the SGRM effects in the indicated assays clustered according to cell type. **(B–C)** LPML correlation plots Pearson correlation, r, calculated from SGRM activity excluding Dex and vehicle. Effects of SGRMs on NTKO cell viability and in the indicated assays in CD8+ Teff cells. Each dot represents a separate compound, with indicated compounds colored as in **Fig. 2**.

**Figure S6.**
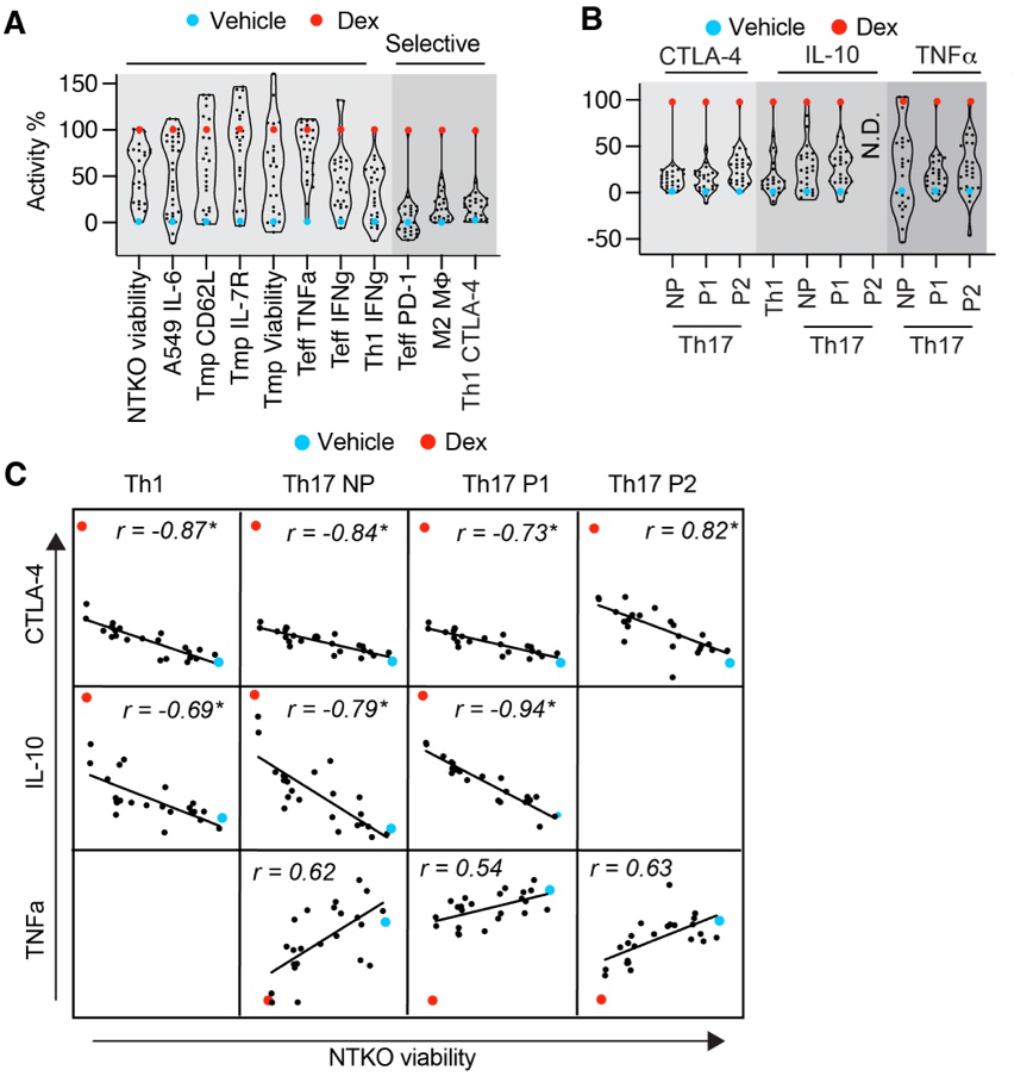
CD4+ cell correlated signaling. **(A-B)** Violin plots displaying the distribution of SGRMs effects as % Dex. The distribution of E_max_ for 31 SGRMs across multiple functional assays is normalized to dexamethasone (red point) and vehicle (blue dot). **(C)** Correlation analysis of SGRM effects on NTKO viability versus expression of markers TNFα, IL-10, and CTLA-4 by Th17 subpopulations, with linear regression lines shown in red.

